# Mutation bias can shape adaptation in large asexual populations experiencing clonal interference

**DOI:** 10.1101/2020.02.17.953265

**Authors:** Kevin Gomez, Jason Bertram, Joanna Masel

## Abstract

The Extended Evolutionary Synthesis invokes a role for development in shaping adaptive evolution, which in population genetics terms corresponds to mutation-biased adaptation. Critics have claimed that clonal interference makes mutation-biased adaptation rare. We consider the behavior of two simultaneously adapting traits, one with larger mutation rate *U*, the other with larger selection coefficient *s*, using asexual traveling wave models. We find that adaptation is dominated by whichever trait has the faster rate of adaptation *v* in isolation, with the other trait subject to evolutionary stalling. Reviewing empirical claims for mutation-biased adaptation, we find that not all occur in the “origin-fixation” regime of population genetics where *v* is only twice as sensitive to *s* as to *U*. In some cases, differences in *U* are at least ten to twelve times larger than differences in *s*, as needed to cause mutation-biased adaptation even in the “multiple mutations” regime. Surprisingly, when *U* > *s* in the “diffusive-mutation” regime, the required sensitivity ratio is also only two, despite pervasive clonal interference. Given two traits with identical *v*, the benefit of having higher *s* is surprisingly small, occurring largely when one trait is at the boundary between the origin-fixation and multiple mutations regimes.

## 1 Introduction

What shapes the course of adaptive evolution? The neo-Darwinian position was that natural selection is preeminent; variation is plentiful and appears in gradual increments, providing raw material that natural selection can shape in any way. The idea that biases in the introduction of variation had a significant influence on the course of evolution was mocked as “the revolt of the clay against the power of the potter” [1]. This selectionist view persisted through the Modern Synthesis [2], following the metaphor of populations as vast “gene pools” with ample amounts of genetic variation already available for natural selection to act on. With abundant mutations each of tiny effect already present and available to recombine to produce any possible phenotype, bias in variation was thought irrelevant.

While the influence of mutation bias on neutral evolution was later acknowledged as part of Neutral Theory, selectionism with respect to the course of adaptation is still upheld by advocates for “Standard Evolutionary Theory”, on the grounds that other evolutionary processes like mutation, drift, and migration, are random with respect to the adaptive direction favored by selection [3]. In contrast, advocates for an “Extended Evolutionary Synthesis” invoke a significant role for developmental processes [4, 5, 6], which shape the phenotypic effects that mutations are more or less likely to have. Exchanges between the two camps have been heated [5, 7].

A role for differences in beneficial mutation rates in shaping the nature of adaptations that evolve has been called “survival of the likeliest” [8], or “first-come-first-served” [9]. Here we refer to it as mutation-biased adaptation. This phenomenon now has substantial empirical support [10]. Evidence for adaptation aligned with mutation bias has been found during the experimental evolution of microvirid bacteriophage [11, 12], *Escherichia coli* [13], and *Pseudomonas aeruginosa* [14, 15]. Studies of parallel adaptation in natural populations also reveal patterns suggestive of mutation bias, e.g. in sodium pump ATP*α*1 adaptations enabling the consumption of glycosine toxins by insects [15], antibiotic resistance in *Mycobacterium tuberculosis* [9], and hemoglobin adaptations for high altitude in birds [16].

A formal population genetic description of mutation-biased adaptation was given by Yampolsky and Stoltzfus [17] for the case where the product of the beneficial mutation rate *U* and the population size *N* is small. In this “origin-fixation” or “strong-selection weak-mutation” parameter regime, each adaptive mutation is either lost or fixed before the next appears [17, 8]. Mutations of each kind appear at their own characteristic mutation rate *U*, and fix with probability proportional to the selection coefficient *s* [18]. Differences in mutation rate and differences in selection thus have exactly equal quantitative influence on the flux of adaptive mutations, which is proportional to *UNs*.

Yampolsky and Stoltzfus [17] originally considered a simple sign epistatic landscape of two loci each with two alleles, beginning in a valley, such that mutating to reach one peak precluded reaching the other, and found mutation biased adaptation even when simulating scenarios with *UN* > 1. However, sign epistasis is not required to argue on the basis of the sensitivities of the flux of adaptive mutants to *U* and to *s* [19]. The theoretical argument for mutation-biased adaptation in this case is, however, strongly dependent on the assumption that the influx of adaptive mutations is small, such that *UN* log(*Ns*) ≪ 1. When two adaptive mutations escape drift to reach appreciable frequencies at the same time, clonal interference will favor the fixation of the one with higher *s* [20]. For this reason, previous theoretical arguments in favor of mutation-biased adaptation have been restricted to the parameter regime *UN* log(*Ns*) ≪ 1, and the phenomenon has been criticized on this basis [3]. However, clonal interference may occur in at least some of empirically documented cases of mutation-biased adaptation [14, 21, 12]. There is therefore a need for theory to understand how mutation-biased adaptation might be possible in the presence of clonal interference.

Models that have previously been used to dismiss mutation-biased adaptation when *UN* log(*Ns*) ≥ 1 [3] are simple, and do not capture the multi-locus complexities included in more recent traveling wave models of population genetics [22, 23, 24, 25]. Here we build on a recent two-dimensional travelling wave model [26] to formally investigate the circumstances under which mutation-biased adaptation can occur.

## 2 Materials and Methods

### (a) Elasticities

We quantify the sensitivity of a population’s rate of adaptation *v* to changes in *U* and *s* by computing elasticities, a metric in common usage in economics (e.g. price elasticity) and applied mathematics, but not in evolutionary biology. Elasticity is the percent change in a function’s output due to a percent change in its input. Given the function *v*(*U, s*), the *U* -elasticity *E*_*U*_ and *s*-elasticity *E*_*s*_ of *v* are 

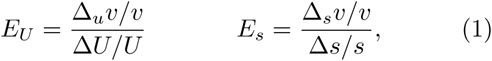

where Δ_*u*_*v* = *v*(*U* + Δ*U, s*) −*v*(*U, s*) and Δ_*s*_*v* = *v*(*U, s* + Δ*s*)*v*(*U, s*). The ratio *E*_*s*_*/E*_*U*_ measures the relative sensitivity of adaptation to selection versus mutation.

The ratio of elasticities *E*_*s*_*/E*_*U*_ can also be calculated using contour lines of *v*(*U, s*), which are curves *U* (*s*) in *U* − *s* space for which *v*(*U* (*s*), *s*) is constant. We show in Section 1 of the Supplement that 

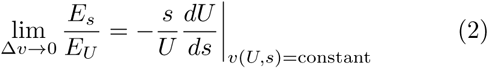

when *U* (*s*) is a contour line of *v*(*U, s*) (see Eq. (S1.1)).

### (b) Origin-fixation regime analytics

Consider the origin-fixation regime (figure 1a), where the rate of adaptation is

**Figure 1:**
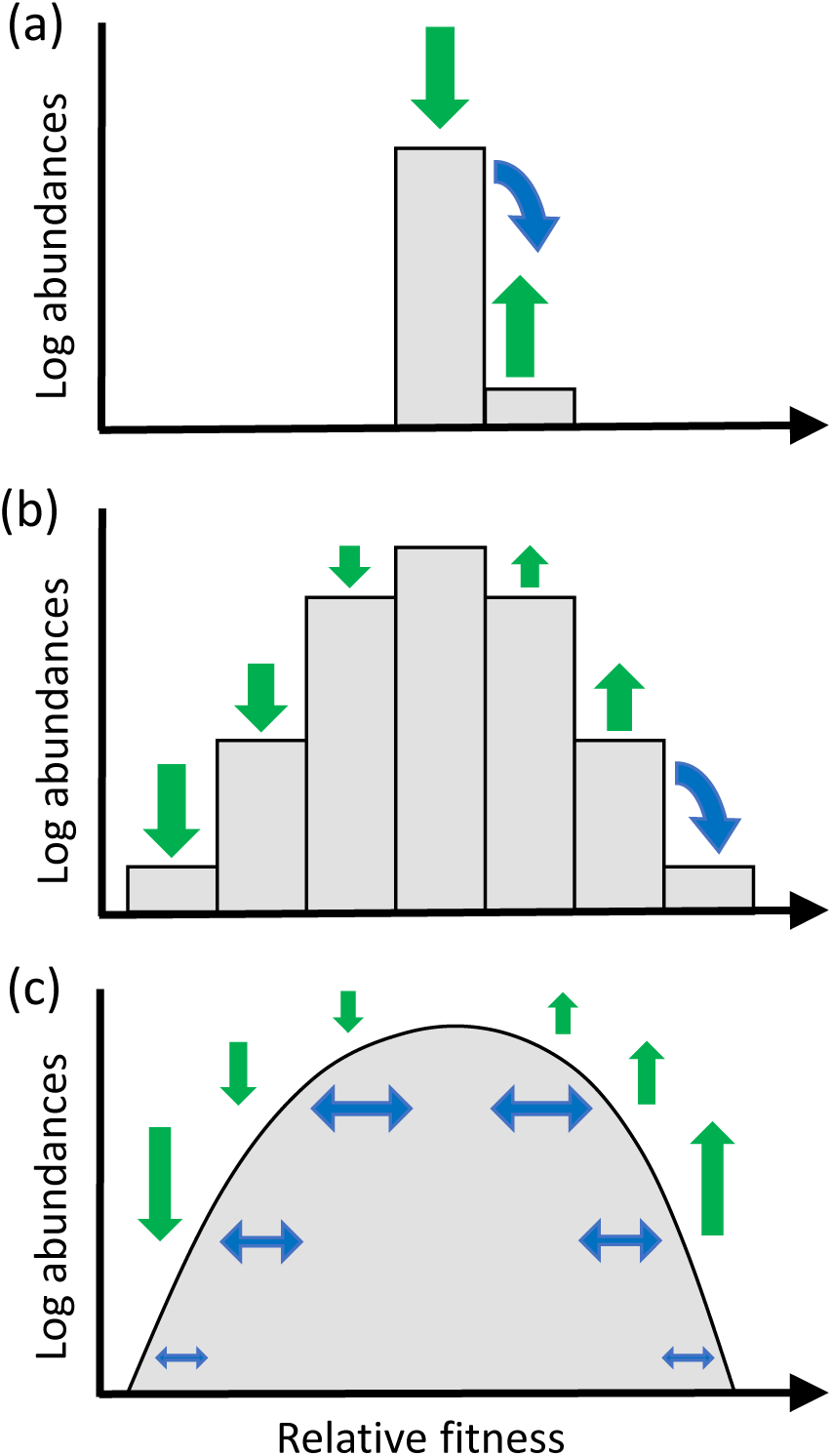
Contribution of mutation fluxes to adaptive evolution in three regimes of adaptation: (a) origin-fixation regime, (b) multiple mutations regime, and (c) diffusive-mutation regime. In all three regimes, the mean fitness moves to the right (adaptation) due to selection. In (a) and (b), changes in fitness class abundances are dominated by selection (green arrows). Mutation fluxes are negligibly small by comparison, except at the front (blue arrow) where they have the crucial role of producing new fitter lineages, allowing the travelling fitness wave to continue. In (c), mutation fluxes when *U ≫ s* increase fitness variance throughout the population (blue double-sided arrow).

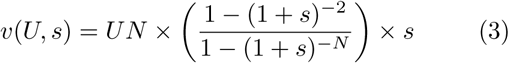

for a population of size *N* .Here *UN* is the total influx of beneficial mutations per generation and [1 − (1 + *s*)^−2^]*/*[1 − (1 + *s*)^−*N*^] ≈ 2*s* is the fixation probability of each mutation [27, Equation 2], with the approximation being valid for *s* ≪ 1. Thus, approximately *UN* × 2*s* mutations are expected to fix in the population each generation, with each increasing relative fitness by *s*, yielding Eq. (3).

The *U* - and *s*-elasticities of *v* in the origin-fixation regime are *E*_*U*_ ≈ 1 and *E*_*s*_ ≈ 2 since Δ_*u*_*v/v* ≈ Δ*U/U* and Δ_*s*_*v/v* ≈ 2Δ*s/s*. The latter expressions follow from applying the first order Taylor approximations 

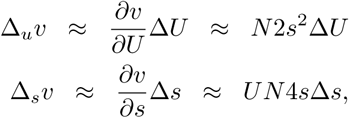

and dividing through by *v* ≈ *UN* 2*s*^2^.

As an alternative way to arrive at the same elasticity ratio, local parameterizations of *v*-contours have the form *U* (*s*) ≈ *v/*2*Ns*^2^, which leads to lim_Δ*v→*0_ *E*_*s*_*/E*_*U*_ = 2 by applying Eq. (2).

This ratio means that in the origin-fixation regime, mutation-biased adaptation will occur when relative differences in *U* (Δ*U/U*) exceed relative differences in *s* (Δ*s/s*) by at least a factor of *E*_*s*_*/E*_*U*_ ≈ 2.

### (c) Multiple mutations regime analytics

When beneficial mutations are more common (*UN* log(*Ns*) ≥ 1), there is competition among lineages, each of which may have accumulated multiple beneficial mutations. The probability of fixation is then no longer a simple function of the selection coefficient of the focal mutation alone, but instead depends on the presence of other beneficial mutations. The resulting complexities change the relationship between *v* and the parameters *N, U* and *s*.

In the staircase model (Desai et al. 28; figure 1b), beneficial mutations appear at rate *U* per birth and each confers a relative fitness advantage of *s* (one “step” on the staircase). The population is divided into fitness classes that are composed of individuals with the same number of (interchangeable) beneficial mutations, and hence the same fitness.

Populations in the multiple mutations regime (*UN* log(*Ns*) ≥ 1) will consist of many fitness classes. A traveling fitness wave results from beneficial mutations adding new fitness classes and selection changing genotype frequencies and eventually culling the least fit classes [23]. A new fitness class appears by mutation at the “nose” of the travelling wave, initially populated by few individuals relative to their selective advantage *Q* over mean population fitness, subject to branching process dynamics dominated by random genetic drift. However, if a new class reaches a size of *n* > 1*/Q*, growth due to selection becomes approximately deterministic. Deterministic classes can be modeled using exponential growth equations, simplifying the analysis of the traveling wave and allowing for efficient numerical simulation as described below in the Subsection Simulations.

Under the assumption that *U* ≪ *s*, Desai and Fisher [23, Eq. 41] obtained

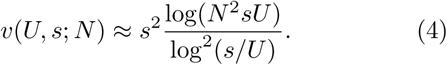

We derive from this the ratio *E*_*s*_*/E*_*U*_ in Supplementary Eq. (S2.5) as 

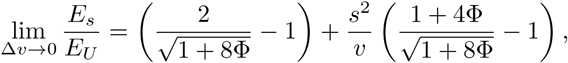

where Φ = *v* log(*Ns*)*/s*^2^. An upper-bound for *E*_*s*_*/E*_*U*_ in the multiple mutations regime is given by Supplementary Eq. (S2.6) 

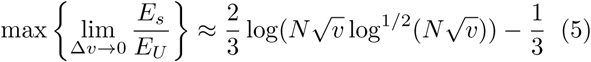

### (d) Diffusive-mutation regime analytics

When *U ≫ s*, the mutational fitness flux between neighbouring fitness classes becomes comparable to the change in fitness class abundance due to selection, breaking down the distinction between deterministic established fitness classes and the stochastic nose. This causes Eq. (4) to break down.

A useful simplifying assumption in this “diffusive-mutation” regime (figure 1c) is to set the beneficial and deleterious mutation rates equal, so that mutation bias or “pressure” [29, 30] alone in the absence of selection does not drive the fitness distribution forward. Mutational steps then follow unbiased random walks with equal chance of reducing or increasing fitness by *s*. The resulting symmetric mutational diffusion process creates fitness variance which selection acts on to drive the wave forward. The diffusion coefficient for this mutational diffusion process is *D* = *Us*^2^*/*2, because the variance in the position of the mutational walk is *s*^2^ per step, and mutations occur at rate *U*. The dependence of *v* on *U* and *s* is then approximately given by [Appendix F in ref 25] 

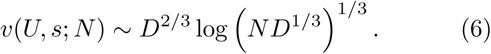

Note that due to the logarithmic second term, the rate of adaptation (variance of the fitness distribution) is approximately proportional to *D*^2*/*3^.

This substantially alters the relative roles of mutation and selection, compared to the multiple mutations regime. From Eq. (6) we calculate the ratio of elasticites to be 

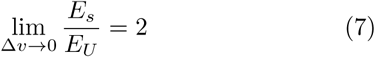

for the diffusive mutations regime (see Supplement, Eq. (S2.8)). This regime could be thought of as a “gene flood” of new mutations, in contrast to the “gene pool” analogy of standing genetic variation in the Modern Synthesis.

### (e) Simulations

The analytical, elasticity-based analysis described above considers evolution in one trait only. But when there is clonal interference, i.e. outside the origin-fixation regime, traits with different *U* and *s* do not evolve independently.

We therefore also simulate the simultaneous evolution of two traits (*k* = 1, 2), with beneficial mutations of fitness effect *s*_*k*_ appearing at rate *U*_*k*_, using an extension of the method of Gomez et al. [26]. We consider strong selection relative to population size, such that 1*/N* ≪ *s*_*k*_ *≤* 1. With no pleiotropy or epistasis in this model, Malthusian fitness is *r*_*i,j*_ = *is*_1_ + *js*_2_ for an individual with *i* beneficial mutations in trait *k* = 1 and *j* beneficial mutations in trait *k* = 2. We group individuals with the same numbers of beneficial mutations into classes denoted by subscripts (*i, j*), with abundances and frequencies at time *t* given by *n*_*i,j*_(*t*) and *p*_*i,j*_(*t*) = *n*_*i,j*_(*t*)*/* ∑_*i′, j′*_ *n*_*i′, j′*_ (*t*), respectively. Thus, the selective advantage of an individual in class (*i, j*) is *Q*_*i,j*_(*t*) = 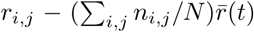 where 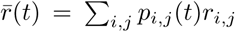 is population mean fitness. Selection according to *Q*_*i,j*_ ensures logistic regulation of population size [31, page 27]. The set of abundances forms a two-dimensional distribution in (*i, j*) space.

Given abundances *n*_*i,j*_(*t*) at generation *t*, we calculate the abundances at generation *t* + 1 in three steps. First, we calculate the expected change due to selection as 

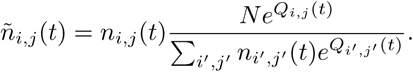

Second, we determine the expected net flux of mutations Δ_*u*_*ñ*_*i,j*_ into class *ñ*_*i,j*_. We also include deleterious mutations in our simulation; these occur at the same mutation rate *U*_*k*_ and decrease fitness by *s*_*k*_, yielding 

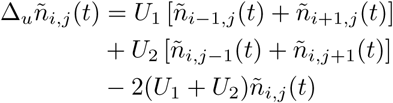

Expected abundances after selection and mutation are 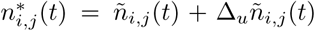. In the last step we set abundances of deterministic classes 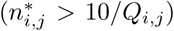 to 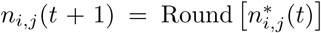, while classes that grow stochastically 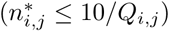 are instead sampled from a Poisson distribution with mean 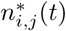.

We implemented the simulation method described above using Matlab code originally developed by Pearce and Fisher [32], modified by Gomez et al. [26] to deal with two traits, and here modified to allow those traits to have distinct mutation rates and selection coefficients. When simulating the evolution of one trait for comparison with theoretical predictions for *v*(*U, s*) given by Eq. (3), Eq. (4) and Eq. (6), we simply set *s*_2_ = 0 and *U*_2_ = 0. We numerically determined *v*(*U, s*) values by applying iterative root-finding methods that allow for noise in *v*(*U, s*) [33]. Time-averages of rates of adaptation were taken over runs lasting 1-2 million generations to ensure convergence, with burn-in periods of 5,000 generations. The length of the burn-in periods is substantially longer than the 1000 generation time to coalescence (sweep time) for all of the parameters sets considered.

Our code ran on Matlab 2016R installed on a desktop computer running Linux. All scripts used to generate our results are available at https://github.com/MaselLab/Gomez_et_al_2020.

## 3 Results

### (a) Visualizing elasticity ratios in the three parameter regimes

To visualize the relative influence of mutation and selection on the rate of adaptation *v*, we plot contour lines of constant *v*(*U, s*) in *U*-*s* parameter space (figure 2), as obtained from analytical theory (Section 2.2; colored piecewise curves) and numerical simulations (colored points). The slopes of these contours as shown in log-log space are closely related to the ratio of elasticities *E*_*s*_*/E*_*U*_, discussed in the Methods as a metric of the relative sensitivities of the adaptation rate to the selection coefficients and appearance rate of beneficial mutations. Thus, steeper contours in figure 2 imply that selection has a greater effect on *v*.

**Figure 2:**
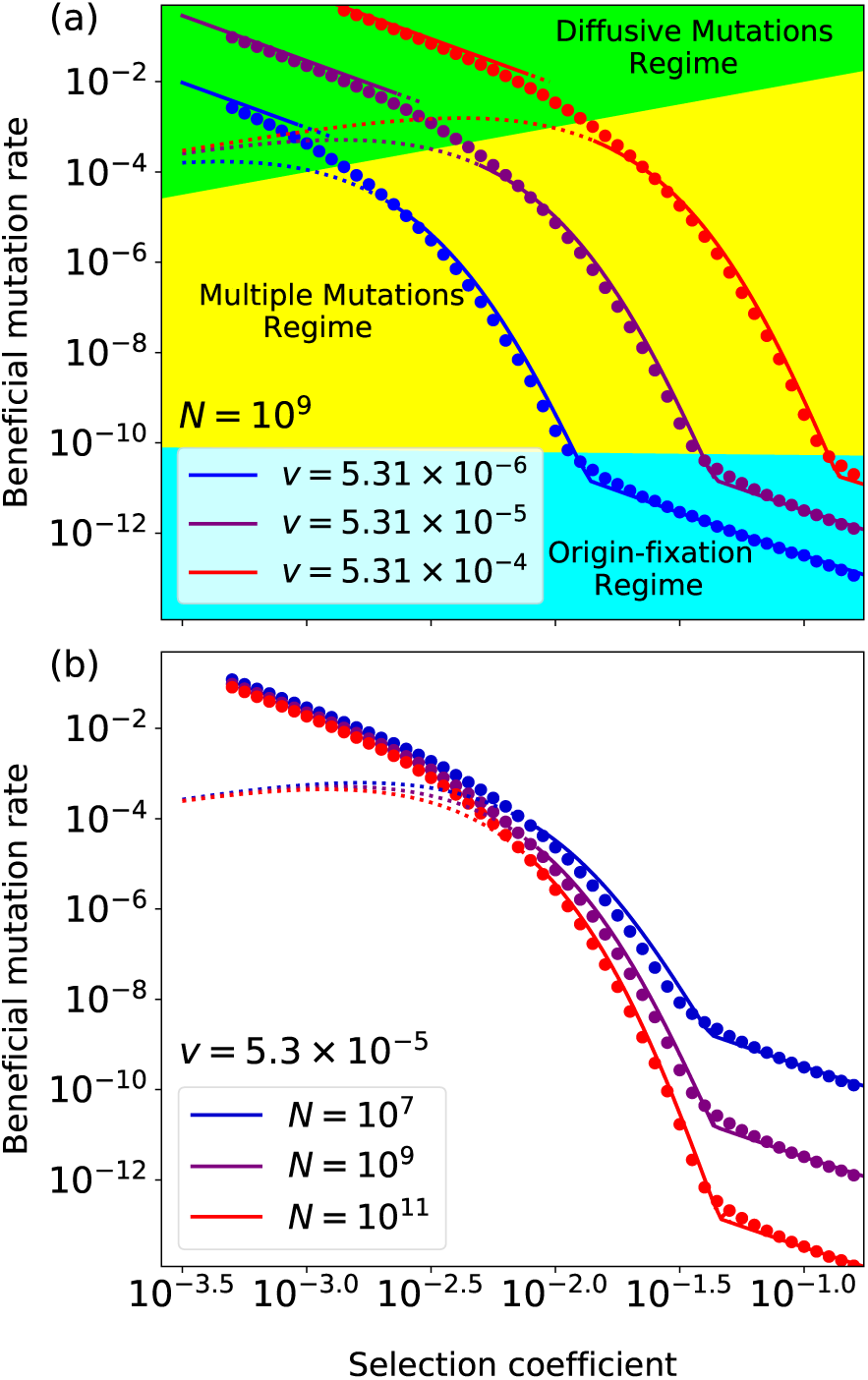
Contour lines of *v*(*U, s*) in *U* -*s* parameter space reveal the possibility of mutation-biased adaptation being just as easy in the diffusive-mutation regime as in the origin-fixation regime. (a) shows distinct *v* contours and (b) shows contours of the same *v* given different values of *N*. Regions of *U* -*s* parameter space in (a) are labeled to indicate where *UN* log(*Ns*) < 1 (origin-fixation; aqua blue), *s* > *U* ≥ 1*/N* log(*Ns*) (multiple mutations; yellow), and *U* ≥ *s* (diffusive-mutation; neon green). In (b), these regime boundaries are not shown since they depend on *N*. Analytic lines for origin-fixation and multiple mutations regimes continue until the point of their intersection. For the transition between multiple mutations and diffusive mutations, where the lines do not cleanly join, we use dotted lines to indicate inappropriate extrapolation beyond the parameter regime of validity.

The slopes of the contours depend on which of the three parameter regimes of *U* and *s* described in the Methods applies, as shown as background color in figure 2a for different values of *v* (figure 2a) and *N* (figure 2b). In the origin-fixation regime (aqua), simulations confirm that *E*_*s*_*/E*_*U*_ ≈ 2, as obtained analytically. As expected from clonal interference, selection becomes more important (*v* contours become much steeper, as high as *E*_*s*_*/E*_*U*_ ≈ 10 − 12) in the multiple mutations regime (yellow region, figure 2a). Figure 2b illustrates how the steepness of *v* contours in the multiple mutations regime changes with population size *N*, as expected from Eq. (5). For sufficiently high mutation rates, the steepness of the contour lines declines and we return to *E*_*s*_*/E*_*U*_ ≈ 2 (green diffusive-mutation region, figure 2a). This implies a shift back to commensurate roles for mutation and selection in adaptation, despite clonal interference.

### (b) Clonal interference creates little bias between two traits of equal *v*

The previous section dealt with the sensitivity of adaptation to *s* and *U* when only a single trait is adapting. But when two traits evolve at the same time, clonal interference may give an advantage to the trait with higher *s*. We therefore simulate the simultaneous evolution of two fitness-associated traits, each of which would on its own evolve at the same rate as the other ({*s,U*} values corresponding to the purple dots in figure 2). We then measured the reduction of a trait’s rate of adaptation due to clonal interference with the second trait. Specifically, we calculated the rate of adaptation for the higher-*s* trait *v*_1_ and divided it by the rate of adaptation of the higher-*U* trait *v*_2_, and examined how this ratio changed across parameter space (figure 3). A ratio higher than one indicates the degree to which clonal interference makes selection more important than indicated by the one-trait elasticity analysis. When both traits are in the origin-fixation regime, we see *v*_1_*/v*_2_ ≈ 1, as expected in the absence of clonal interference (figure 3; pale color in the aqua blue vs. aqua blue lower right).

**Figure 3:**
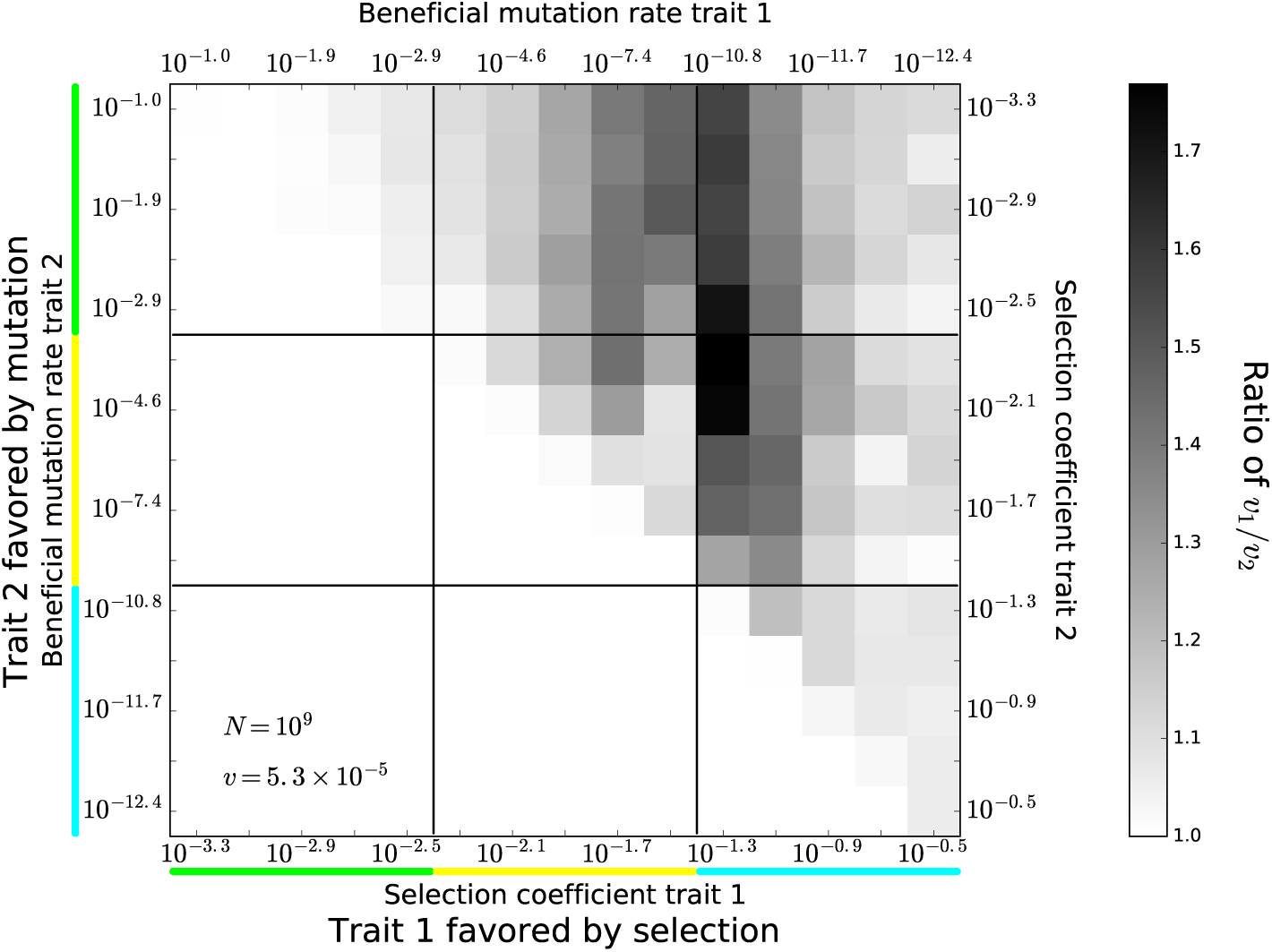
Heat map showing relative rates of adaptation in two simultaneously adapting traits, both of whose values of *U* and *s* would in isolation produce the same rate of adaptation *v* = 5.3 × 10^−5^. Axes are labeled by color to show the regime (from figure 2a) that a trait would be in were it evolving on its own. Darker cell color indicates how much faster the selection-favored trait (*v*_1_) evolves relative to the mutation-favored trait (*v*_2_). Cell values in the lower left are redundant and not shown, as these would amount to switching trait labels. The fact that ratios are near 1 makes stochasticity difficult to eliminate from the visual presentation.

Surprisingly, we see even smaller deviations from *v*_1_*/v*_2_ = 1 when both traits fall within the diffusive-mutation regime (pale color in the neon green vs. neon green upper left of figure 3 never exceeds *v*_1_*/v*_2_ = 1.09). Our finding of *E*_*s*_*/E*_*U*_ ≈ 2 already suggested susceptibility to mutation-biased adaptation; here we see that this persists even when clonal interference is fully accounted for.

Indeed, deviations from *v*_1_*/v*_2_ = 1 are mild throughout, with the highest value of 1.77 observed when the high-*s* trait is near the boundary between origin-fixation and multiple-mutation regimes. At this maximum value, adaptation in the mutation-favored trait still accounts for 36% of the total rate of adaptation. Clonal interference thus has a limited ability to block mutation-driven adaptation, beyond that captured by the sensitivity of *v* to *s*.

### (c) Evolutionary stalling is driven by differences in *v* not *s*

The previous section shows that two traits with the same independent rate of evolution *v*, but drastically different values of *s* and *U*, will continue to adapt at similar rates even in the presence of clonal interference. Next we ask how clonal interference operates between two traits that differ not only in their values of *s* and *U*, but also in their values of *v*. Venkataram et al. [34] demonstrated that functional modules can experience “evolutionary stalling”. Our aim is to explore whether stalling depends only on immediate clonal interference and hence only the relative values of *s*, or whether relative values of *U* also matter.

Before doing so, we note that both *s* and *U* change during adaptation due to diminishing returns epistasis [35] and to the depletion of beneficial mutations, respectively. Future work to explore the interplay between evolutionary stalling and diminishing returns epistasis may build on a recent traveling wave framework capturing absolute fitness [36]. For now we assume fixed values of *s* and *U* associated with each trait.

To test whether evolutionary stalling occurs, and if so whether the identity of the dominant trait depends on *s* alone, we quantify how a focal trait’s adaptation suffers from clonal interference from a second trait, as a function of that second trait’s value of *s* and *U* (figure 4). The top, middle, and bottom panels of figure 4 show focal traits with the same *v* taken from the origin-fixation, multiple-mutations, and diffusive-mutation regimes, respectively (open circles). These three panels are striking in their similarity. Adaptation in the focal trait is essentially unaffected by clonal interference from a trait with smaller *v*, but severely affected by a trait with larger *v*. In other words, we see marked evolutionary stalling that depends on *v*.

**Figure 4:**
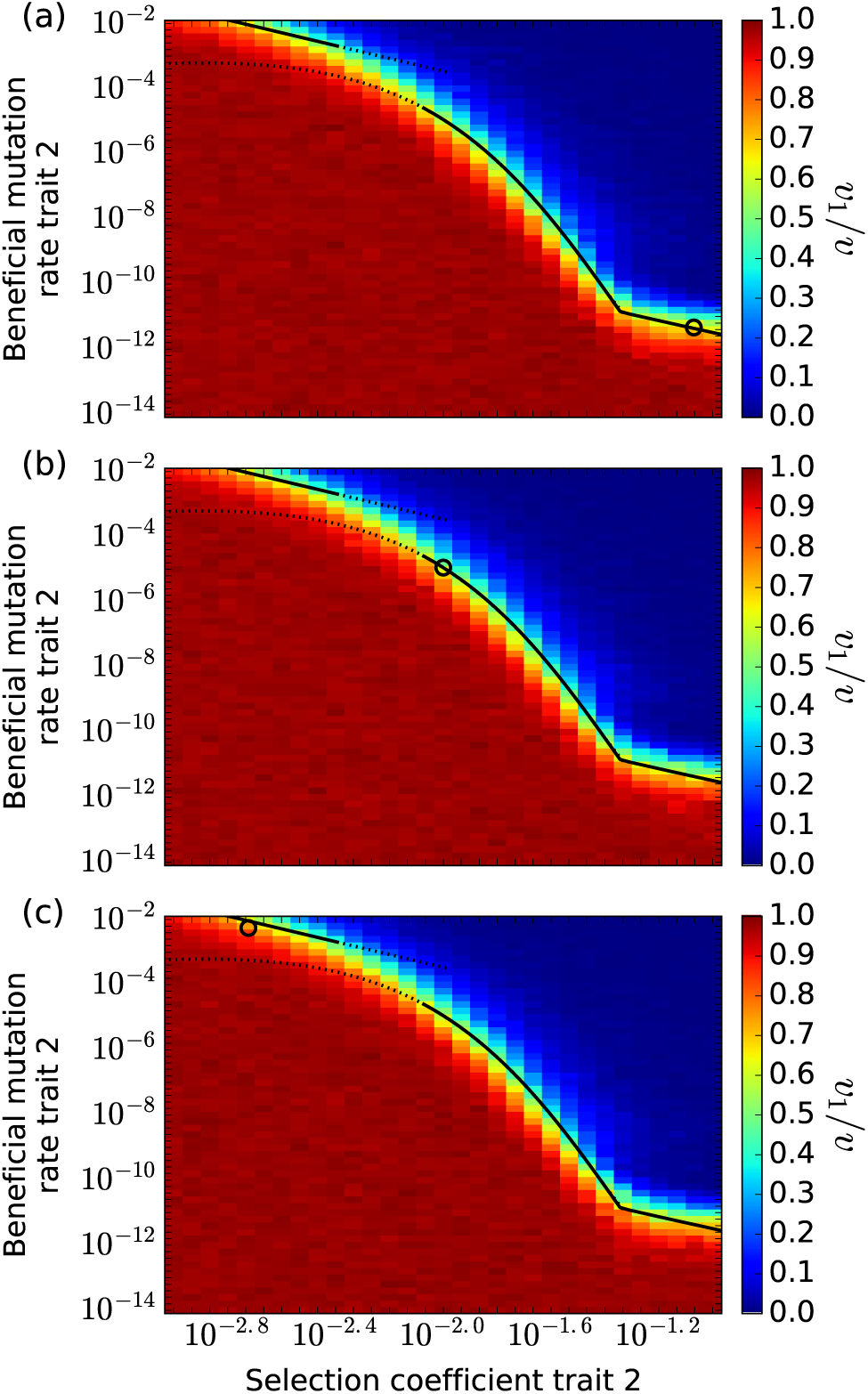
The trait with larger *v* is little affected by clonal interference with another trait, while that with smaller *v* stalls. The focal trait, shown as a dot, would have the same value of *v*_1_ = 5.31 × 10^−5^ in each of the three panels, if it were evolving on its own. The focal trait falls into the origin-fixation regime, multiple mutations regime, and diffusive-mutation regime in the top, middle, and bottom panel, respectively. Colors show the extent to which adaptation in the focal trait slows down as a function of the selection coefficient *s*_2_ and mutation rate *U*_2_ of a second clonally interfering trait. The theoretical *v* contour for *v* = 5.31 × 10^−5^ from figure 2a is shown as a black piecewise curve in each panel. *v*_1_ transitions from a negligible impact of clonal interference from trait two (dark red region) to having its evolution nearly halted by evolution in trait two (dark blue region). The decline in *v*_1_ follows patterns set by *v*-contours, not *s*_2_ or *U*_2_, indicating that the rate of adaptation of trait two, when evolving independently, determines the decline in *v*_1_ from clonal interference between traits. In all panels *N* = 10^9^.

We analytically solve for vertical cross-sections of figure 4 in Section 3 of the Supplement. Eq. (S3.12) gives the color scale along the cross-section as 

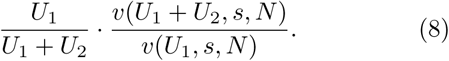

The first term in Eq. (8) drives strong stalling once *U*_2_ *≫ U*_1_.

### (d) Empirical examples of mutation-driven adaptation include the multiple mutations regime

Table 1 summarizes a variety of proposed empirical examples of mutation-biased adaptation. In Table 1 we attempt to quantitatively estimate *N, s* and *U*. When this is not possible, we qualitatively assess which parameter value regime likely applies. If the evidence for mutation-biased adaptation is strong, this can sometimes provide some information about what the missing parameter values must be, to be compatible with this evidence.

**Table 1:** Empirical examples of mutation-biased adaptation. In Couce et al. [13], antibiotic concentrations were titrated to allow proliferation of *E. coli*. The beneficial mutation rate for [13] was assumed to be 3% of the total point mutation rate in the mutator strain studied. The population sizes reported for Schober et al. [43] and Rodrigues and Shakhnovich [42] are CFU values calculated from the optical densities (OD_600_) reported in their papers. We used the formula CFU = OD_600_ · (8 × 10^8^) · volume, where volume refers to culture volume (ml) in a turbidostat [43], or fresh media volume in 96-well flat-bottom plates [42]. The factor 8 × 10^8^ is the scaling constant yielding CFU/ml = OD_600_ · 8 × 10^8^. Entries in which quantitative ranges for *N, s* and *U* are not known are assigned a qualitative description (large, small, tiny, etc.) to indicate the logic by which we used their relative sizes to infer which parameter regime applies. Stoltzfus and McCandlish’s [15] meta-analysis, which examines several natural populations and evolution experiments, is designated as a mixture of origin-fixation and multiple mutations regimes.

| System/Organism | Natural / Lab | Population size ($N$) | Selection coeff. ($s$) | Beneficial mut. rate ($U$) | Regime | Estimated mutation bias | Comment |
| --- | --- | --- | --- | --- | --- | --- | --- |
| 70 avian taxa; Hb-O2 affinity adaptations to elevation<br>Storz et al. [16] | natural | small to moderate | moderate to large | small because few genes | OF | 6 fold for mutations | Excess of CpG adaptive mutations compared to number predicted by null model where CpG status irrelevant to adaptation |
| De novo gene birth in overlapping reading frame in 80 capsid viral species<br>Willis & Masel [38] | natural | large | unknown | tiny | OF because tiny $U$ | $> 465:92 = 5$ | Mutation bias greater if longer ORFs are more likely to be adaptive |
| Meta-analysis of adaptive mutations<br>Stoltzfus & McCandlish [15] | natural & lab | mixed | mixed | mixed | Mixed, OF + MM | 3 – 7 fold for mutations,<br>2 – 4 fold for unique mutations | Excess transitions |
| *M. tuberculosis*; antibiotic resistance<br>Payne et al. [9] | natural clinical isolates | large | large<br>100 fold<br>increase in MIC | small<br>$10^{-9} - 10^{-7}$ | OF or MM | 1.6 – 41.6 fold for mutations | Excess transitions for 9 out of 11 different antibiotics |
| *P. aeruginosa*; antibiotic resistance<br>MacLean et al. [14] | lab | $10^4 - 10^9$ | 0.12<br>– 0.96 | $8.3 \times 10^{-10}$<br>– $2.5 \times 10^{-8}$ | OF or MM | Allele-specific mutation rate varies by 30 fold | In 11 adaptive mutations, chances of evolving correlated with respective mutation rates [10], and uncorrelated with selection coefficients |
| ssDNA bacteriophages ID8, ID11, NC13, WA13; phage growth<br>Sackman et al. [12] | lab | $10^5 - 10^9$ | 0.11<br>– 0.64 | $1.1 \times 10^{-3} - 5.3 \times 10^{-3}$<br>(total mut. rate) | OF or MM | 24 fold for mutations,<br>11 fold for unique mutations | Excess transitions |
| ssDNA bacteriophage ID11; phage growth<br>Rokyta et al. [11] | lab | $10^4 - 10^8$ | 0.11<br>– 0.39 | $1.1 \times 10^{-3} - 5.3 \times 10^{-3}$<br>(total mut. rate) | OF or MM | 8 – 9 fold among relative mutation rates | OF model used to estimate relative mutation rates. Ranking of $s$ differs from [12]. |
| HIV-1; evolved in cultures of human T-cells<br>Bertels et al. [37] | lab | $1 \times 10^3 - 6 \times 10^6$ | large | high | MM | 5 fold for mutations,<br>6 fold for unique mutations | Excess transitions |
| *E. coli*; evolved with DHFR inhibited using antibiotic<br>Schober et al. [43] | lab | $6 \times 10^6 - 7.2 \times 10^8$ | large | high | MM | unavailable | Mutations rewiring broken metabolic network to less efficient configuration favored over those restoring original efficient configuration |
| *E. coli*; adaptation following inactivation of DHFR<br>Rodrigues et al. [42] | lab | $1.6 \times 10^7 - 4.8 \times 10^7$ | large | small | MM | unavailable | Mutations rewiring broken metabolic network to less efficient configuration favored over those restoring original efficient configuration |
| *E. coli*; antibiotic resistance<br>Couce et al. [13] | lab | $7.2 \times 10^7 - 3.6 \times 10^9$ | large<br>0.00158<br>– 1 | high<br>$3 \times 10^{-4}$<br>– $1 \times 10^{-2}$ | MM or DM | 100 – 300 fold ($\Delta mutH$),<br>500 – 10,000 fold ($\Delta mutT$) | Different mutational profiles of $\Delta mutT$ (elevates transversions, excl. usual transition adaptations) v. $\Delta mutH$ (elevates transitions) |

Note that many but not all of these empirical examples argue on the basis transition : transversion ratios [15, 9, 12, 37]. According to a selection-driven model, in which each mutation is equally likely to be beneficial, adaptive point mutations should have a 1 transition : 2 transversion ratio. However, the mutational spectrum is biased toward transitions, and so an excess of transitions is evidence for mutation-biased adaptation.

The first two entries in Table 1 are expected to lie in the origin-fixation regime, albeit for different reasons. *N* is likely reasonably small for the natural bird populations studied by Storz et al. [16]. While *N* is much larger for the natural viral populations studied by Willis and Masel [38], the rate *U* of the beneficial *de novo* birth of new overlapping genes is likely so low that *UN* ≪ 1. There were 5 times more birth events in the +1 reading frame, which has more and longer ORFs, than there were in the +2 reading frame, on which the genetic code bestows a tendency toward the favored protein property of high intrinsic structural disorder. Neither study quantified differences in *U* or *s* — instead, differences in occurrence, i.e. in *v/s*, were counted.

In a meta-analysis of 15 studies (10 natural and 5 experimental) of different taxonomic groups, potentially representing a mixture of origin-fixation and multiple mutations regimes, Stoltzfus and McCandlish [15] detected a 3-fold excess of transitions among all natural adaptive point mutations, representing a 2-fold excess of unique adaptive mutations relative to the expected 1:2 ratio. In experimental cases, 7-fold and 4-fold excesses, were seen for mutations and unique mutations, respectively.

All the remaining entries in Table 1 involve microbial experimental evolution, which generally takes place in the multiple mutations regime [39, 40, 41], although we are not able in all cases to rule out the possibility that bottlenecks create more of an origin-fixation regime. Given that these studies report large differences between mutation rates, compatible with *E*_*s*_*/E*_*U*_ in the range between 2 and 10, mutation-biased adaptation can take place even in the multiple mutations regime.

Both Rodrigues and Shakhnovich [42] and Schober et al. [43] examined experimental evolution to recover from a genetic loss, and found it to be mutationally easier to inactivate more loci, rewiring metabolic pathways in the process, than to recover the efficiency of the original pathway. This is mutation-biased adaptation to the detriment of long-term adaptation.

In contrast, Venkataram et al. [34] partly disabled the translational machinery and then observed this previously highly conserved complex actively evolve to recover. Then as adaptive improvements accumulated, diminishing returns epistasis led to a drop in *s*, and evolution again “stalled”. Our theoretical results support the concept that evolution of a module will stall once its value of *v* in isolation is no longer higher than that of other modules.

The only candidate for the diffusive-mutation regime in Table 1 is Couce et al.’s [13] experiments with two mutator strains of *E. coli*, i.e. with artificially high *U*. One mutator strain (ΔmutH) has transition rates 100-300 times larger than wild-type [44, 45], the other (ΔmutT) has transversion rates 500-10,000 times larger than wild-type [44, 46]. These high mutation rates make *U* > *s* and hence the diffusive-mutation regime a possibility, although not certain; examples of adaptation in the diffusive-mutations regime typically invoke RNA viruses [47].

These empirical examples of mutation-biased adaptation collectively span two or even three regimes of adaptation, and in particular, include evolution experiments known to include rampant clonal interference in the multiple mutations regime. Mutation-biased adaptation in this regime requires large mutation bias relative to selective differences. Unsurprisingly, where relative mutation rates have been quantified, biases of appropriate magnitude were found. We can therefore infer in cases where relative mutation rates are not known, but mutation-biased adaptation has been documented in the multiple mutations regime, that similarly large mutation bias must exist. Thus, one reason that mutation-biased adaptation may be more widespread than previously claimed [3] is that differences in *U* can be much larger than differences in *s*.

## 4 Discussion

When two adaptive traits evolve together, adaptation is dominated by whichever has the higher rate of adaptation *v* in isolation, not by whichever has the higher selection coefficient *s*. If differences in *v* among fitness-associated traits were dominated by differences in *s*, then this would be a distinction without a difference. But if differences in *U* are sufficiently large relative to differences in *s*, then mutation-biased adaptation will occur. How much larger differences in *U* need to be is well summarized using simple equations for the adaptation rate *v*, and depends on which parameter value regime the population is in. For both the origin-fixation regime with *UN* log(*Ns*) ≪ 1, and the diffusive-mutation regime with *U* > *s*, differences in *U* need to be twice as large as differences in *s*. For values of *U* in the multiple mutations regime between the two, ratios of as much as 10-12 might be required. Our theoretical calculations in the multiple mutations and diffusive-mutation regime are for the strong linkage disequilibrium produced in asexual microbes — in the absence of strong linkage disequilibrium in more sexual populations, the origin-fixation regime is likely a good approximation.

### (a) Why does *s* not matter beyond *v*?

Because clonal interference between two genotypes depends on *s*, it is at first unintuitive that stalling depends on *v* rather than *s*. To understand why, consider adaptation with one trait in the multiple mutations regime and a second in the origin-fixation regime. Because mutations are rare in the second trait, clonal interference between traits is minimal. However, each selective sweep of a beneficial mutation in trait two (origin-fixation) decreases genetic variation in trait one (multiple mutations). When the *v* of trait two is larger, these sweeps occur more frequently and lead to fewer substitutions in the first trait, further stalling its evolution.

Previous work has conjectured that adaptation might be dominated by beneficial mutations with the largest *v* [48, 23, 49]. Here we have shown that this conjecture was correct (figure 4).

### (b) Deleterious mutations

For mathematical convenience in the diffusive-mutation regime, we assumed throughout that the deleterious mutation rate is equal to the beneficial mutation rate. In fact, deleterious mutations are much more common. Representing this asymmetry would have no effect on the dynamics of adaptation in the origin-fixation regime. In the other two regimes, assuming symmetry in effect sizes, larger deleterious mutation rates would primarily alter the shape of the traveling wave [23, 50, 51], rather than alter *v*. Deleterious mutations do contribute to *v* when their effect sizes are smaller, but do so in a linear manner, resulting in *v* = *v*_*b*_ − *v*_*d*_ [51]. Our analysis then has its basis in (*v*_*b*_), such that the elasticities we calculate should remain valid.

### (c) Conclusion

Both selective advantages *s* and beneficial mutation rates *U* determine adaptation rates, which in turn determine which trait will dominate the adaptive process. Differences in *U* need to be 2 to 10 times as large, depending on the parameter value regime, in order to swamp differences in *s*. Both molecular and developmental biases can create such large differences in *U*, leading to mutation-biased adaptation. Diverse case studies suggest that mutation bias significantly shapes which adaptations occur, even in populations with strong clonal interference. While adaptation does not occur without natural selection, which adaptation occurs among the many possibilities has more complex causes.

## Acknowledgments

We thank Benjamin Good for helpful discussions and Arlin Stolzfus and Sergey Kryazhimskiy for comments on the manuscript. KG, JB, and JM were funded by the National Science Foundation (DEB-1348262), KG by the National Institutes of Health (T32 GM084905) and JB by the Environmental Resilience Institute at Indiana University.

## Notes

https://github.com/MaselLab/Gomez_et_al_2020

### Competing Interest Statement

The authors have declared no competing interest.

### Summary of Updates

Methods and results reorganized, and new supplement added. Minor revisions were also made to the manuscript's contents regarding the discussion of elasticities.

