## Supplement for "Mutation bias can shape adaptation in large asexual populations experiencing clonal interference"

#### 1 Calculating $E_s/E_U$ from the slopes of $v$ contours

Here we derive the relationship between slopes of $v$ contours and the ratio of elasticities $E_s/E_U$. In particular, we will show that

$$\lim_{\Delta v \to 0} \frac{E_s}{E_U} = -\frac{s}{U(s)}\frac{dU(s)}{ds}. \tag{S1.1}$$

when $U(s)$ is a parameterization of a $v$ contour. Eq. (S1.1) will be used to calculate the ratio of elasticities $E_s/E_U$ across the various regimes.

Consider a solution $U(s)$ to the equation $v(U(s), s) = v_c$ ($v_c$ constant), where Eq. (2), Eq. (3), and Eq. (5) give $v(U, s)$ in their respective regimes. Implicit differentiation of both sides of $v(U(s), s) = v_c$ with respect to $s$, and application of the chain rule provides

$$\frac{\partial v}{\partial U}\frac{dU}{ds} + \frac{\partial v}{\partial s} = 0.$$

Multiplying this expression by $\frac{(\Delta s)^2}{Uv}\frac{dU}{ds}$ gives

$$0 = \left(\frac{\Delta s}{v}\frac{\partial v}{\partial U}\right)\left(\frac{\Delta s}{U}\frac{dU}{ds}\right) + \left(\frac{\Delta s}{v}\frac{\partial v}{\partial s}\right)\left(\frac{\Delta s}{s}\right)\left(\frac{s}{U}\frac{dU}{ds}\right). \tag{S1.2}$$

For small changes in the selection coefficient $\Delta s$, $\frac{\Delta U}{U} \approx \frac{\Delta s}{U}\frac{dU}{ds}$, and consequently

$$\frac{\Delta_u v}{v} \approx \frac{\Delta s}{v}\frac{\partial v}{\partial U} \qquad \text{and} \qquad \frac{\Delta_s v}{v} \approx \frac{\Delta s}{v}\frac{\partial v}{\partial s}.$$

In addition,

$$\frac{d(\log U)}{d(\log s)} = \frac{s}{U}\frac{dU}{ds} \tag{S1.3}$$

also holds, where $\frac{d(\log U)}{d(\log s)}$ is the slope of the $v$ contour in the $\log(U) - \log(s)$ plane. Substituting for these terms in Eq. (S1.2) and rearranging provides

$$-\frac{d(\log U)}{d(\log s)} = \lim_{\Delta v \to 0} \frac{\left(\frac{\Delta_s v}{v}\right) / \left(\frac{\Delta s}{s}\right)}{\left(\frac{\Delta_u v}{v}\right) / \left(\frac{\Delta U}{U}\right)}.$$

The left side is the slope of a $v$ contour line in $\log U$ vs. $\log s$ space, scaled by $-1$ and evaluated at $s$, while the right side is the ratio $E_s/E_U$ in Eq. (S1.1). This establishes Eq. (S1.1). Note that we have used the fact that for small changes in a quantity, relative changes are approximately equal to changes in the logarithm of that quantity, eg. $\Delta s/s \approx \Delta \log s$.

#### 2 Parameterizing $v$-contours in the multiple mutations and diffusive mutations regimes

We first derive parameterizations of $v$ contour lines in the multiple mutations regime analyzed by Desai and Fisher [1]. Eq. (4) for $v(U, s)$ can be rewritten as

$$\ell^2 + \frac{s^2}{v}\ell - \frac{2v\log(Ns)}{s^2} = 0.$$

where $\ell = \log(s/U)$. The quadratic polynomial in $\ell$ has solutions

$$\ell = \frac{s^2}{2v}\left(\pm\sqrt{1+\frac{8v\log(Ns)}{s^2}} - 1\right).$$

However, the positive root is the correct solution since $\ell = \log(s/U) > 0$ when $U/s < 1$, as assumed in [1] and distinguishing the multiple mutations regime from the diffusive mutations regime. Solving for $U$ in

$$\log(s/U) = \frac{s^2}{2v}\left(\sqrt{1+\frac{8v\log(Ns)}{s^2}} - 1\right)$$

provides the parameterization

$$U(s) = s\exp\left(-\frac{s^2}{2v}\left(\sqrt{1+\frac{8v\log(Ns)}{s^2}} - 1\right)\right). \tag{S2.4}$$

Equation (S2.4) allows us to calculate $E_s/E_U$ in the multiple mutations regime by applying Eq. (S1.1). In particular, we have

$$\begin{aligned}
\lim_{\Delta v\to 0}\frac{E_s}{E_U} &= -\frac{s}{U(s)}\frac{dU(s)}{ds} \\
&= -\frac{s}{U(s)}\left[\frac{U(s)}{s} + U(s)\cdot\frac{d}{ds}\left(-\frac{s^2}{2v}\left(\sqrt{1+\frac{8v\log(Ns)}{s^2}} - 1\right)\right)\right] \\
&= -1 + s\left[\frac{s}{v}\left(\sqrt{1+\frac{8v\log(Ns)}{s^2}} - 1\right) + \frac{2}{s}\left(\frac{1-2\log(Ns)}{\sqrt{1+\frac{8v\log(Ns)}{s^2}}}\right)\right] \\
&= -1 + \frac{s^2}{v}\left(\sqrt{1+\frac{8v\log(Ns)}{s^2}} - 1\right) + 2\left(\frac{1-\frac{2s^2}{v}\cdot\frac{v\log(Ns)}{s^2}}{\sqrt{1+\frac{8v\log(Ns)}{s^2}}}\right)
\end{aligned}$$

which simplifies to

$$\lim_{\Delta v\to 0}\frac{E_s}{E_U} = \left(\frac{2}{\sqrt{1+8\Phi}} - 1\right) + \frac{s^2}{v}\left(\frac{1+4\Phi}{\sqrt{1+8\Phi}} - 1\right), \tag{S2.5}$$

where $\Phi = v\log(Ns)/s^2$. Eq. (S2.5) provides the value of $E_s/E_U$ as a function of $s$ in the multiple mutations regime.

The maximum value of $E_s/E_U$ occurs near the transition between the multiple mutations regime and the origin-fixation regime. To calculate this value, we note that in the origin-fixation regime, $v \approx 2NUs^2$ and $NU\log(Ns) \leq 1$. Thus, $U = v/2Ns^2$ and $\Phi = v\log(Ns)/s^2 \leq 1$ for a fixed value of $v$. The transition between the regimes occurs at $s_*$ where $\Phi = v\log(Ns_*)/s_*^2 \approx 1$. We can obtain an approximate solution for $s_*$ by solving the equivalent equation

$$s_* = f(s_*) = \sqrt{v}\log(Ns_*)^{1/2}.$$

The function $f(s) = \sqrt{v}\log(Ns)^{1/2}$ satisfies the properties $f(I) \subset I$ and $\max_{s\in I}|f'(s)| < 1$ on the closed interval $I = [\sqrt{v}, 1]$ when $\log(N\sqrt{v}) > 1$ and $\sqrt{v}\log(N)^{1/2} < 1$. Thus, $f(s)$ is a contraction mapping on $I$ with fixed point solution $s^* \in I$. We define the sequence $\{s_n\}_{n=1}^{\infty}$ with $s_0 = 1$ and $s_n = f(s_{n-1})$ for $n \geq 1$. The *contraction mapping theorem* implies that $\lim_{n\to\infty} s_n = s_*$ [2, see p.220]. We select the endpoint $s_0 = 1$ to obtain faster convergence in $\{s_n\}_{n=1}^{\infty}$ since $\max_{s\in[s_*,1]}|f'(s)| < \sqrt{v}$. Two iterations of the sequence provides the estimate

$$s_* \approx s_2 = f(f(1)) = \sqrt{v}\log(N\sqrt{v}\log(N\sqrt{v})^{1/2})^{1/2}.$$

Substituting in $s_*$ and $\Phi = v\log(Ns_*)/s_*^2 \approx 1$ into the left-hand side of Eq. (S2.5) yields

$$\max_{s\in MM}\left\{\lim_{\Delta v\to 0}\frac{E_s}{E_U}\right\} \approx \frac{2}{3}\log(N\sqrt{v}\log(N\sqrt{v})^{1/2}) - \frac{1}{3}, \tag{S2.6}$$

as the maximum value of $E_s/E_U$ in the multiple mutations regime ($s \in MM$). The approximation above indicates that at the onset of clonal interference, the relative importance of selection for adaptation grows stronger with larger population size, as can be seen in Figure 2b.

To obtain a parameterization for contours of $v(U, s)$ given by Eq. (6) for the diffusive mutations regime, we can solve for $U$ in an equivalent equation

$$v \sim \left(\frac{Us^2}{2}\right)^{2/3}\log^{1/3}\left(N\left(\frac{Us^2}{2}\right)^{1/3}\right) \qquad \Longrightarrow \qquad \frac{24v^3}{U^2s^4} \sim \log\left(\frac{6N^6v^3}{\frac{24v^3}{U^2s^4}}\right).$$

This leads to an expression of the form $w = \log\left(6v^3N^6/w\right)$, where $w = 24v^3/U^2s^4$. The function $g(w) = \log\left(6v^3N^6/w\right)$ is a contraction mapping over the bounded interval $I = [1, 6v^3N^6e]$ when $\log(\log(6v^3N^6)) > 1$, so we can use the previous approach discussed above to obtain an approximate solution for $w^*$. We define a sequence $\{w_n\}_{n=1}^{\infty}$ with $w_0 = 1$ and $w_n = g(w_{n-1})$. Then

$$w_* \approx w_2 = g(g(1)) = \log\left(\frac{6v^3N^6}{\log(6v^3N^6/\log(6v^3N^6))}\right).$$

We can then solve for $U$ in

$$\frac{24v^3}{U^2s^4} = w_* = \log\left(\frac{6v^3N^6}{\log(6v^3N^6/\log(6v^3N^6))}\right),$$

which leads to

$$U(s) = \frac{1}{s^2}\left(\frac{2v^{3/2}}{\log\left(N\sqrt{v}/\log\left(N\sqrt[6]{6v^3}\right)\right)^{1/2}}\right). \tag{S2.7}$$

Eq. (S2.7) allows us calculate the $E_s/E_U$ in the diffusive mutations regime by applying Eq. (S1.1). In particular, we have

$$\lim_{\Delta v\to 0}\frac{E_s}{E_U} = -\frac{s}{U(s)}\frac{dU(s)}{ds}$$

$$= -s^3\left(\frac{\log\left(N\sqrt{v}/\log\left(N\sqrt[6]{6v^3}\right)\right)^{1/2}}{2v^{3/2}}\right)\left[-\frac{2}{s^3}\left(\frac{2v^{3/2}}{\log\left(N\sqrt{v}/\log\left(N\sqrt[6]{6v^3}\right)\right)^{1/2}}\right)\right],$$

and thus,

$$\lim_{\Delta v \to 0} \frac{E_s}{E_U} = 2 \tag{S2.8}$$

in the diffusive mutations regime.

### 3 Expected rates of adaptation for two clonally interfering traits in the multiple mutations regime with distinct mutation rates

We derive solutions for $v_i$ $(i = 1, 2)$ of two clonally interfering traits in the multiple mutations regime when the traits have identical selection coefficients, but distinct mutation rates $U_i$. The derivation follows similar steps to those given in Appendix B of Gomez et al. [3], who treated the case where both $s$ and $U$ are identical across the two traits.

We begin by defining an indicator variable $I_{1,i}(l)$ that identifies whether a beneficial mutation $l$ in individual $i$ occurred in trait one. If we consider two individuals, $i$ and $j$, chosen at random from the population, they will each differ by $n_i$ and $n_j$ beneficial mutations from their common ancestor. The mean genetic variance in trait one can then be expressed as

$$\sigma_1^2 = \frac{s^2}{2} E\left[\left(\sum_{l=1}^{n_i} I_{1,i}(l) - \sum_{m=1}^{n_j} I_{1,j}(m)\right)^2\right],$$

where we use the property $\text{var}(X-Y) = 2\text{var}(X)$ when $X$ and $Y$ are independent and identically distributed. Expanding the expression provides

$$\sigma_1^2 = \frac{s^2}{2} E\left[\left(\sum_{l=1}^{n_i} I_{1,i}(l) - \sum_{l=1}^{n_j} I_{1,j}(l)\right)^2\right]$$

$$= \frac{s^2}{2}\left(E\left[\sum_{l=1}^{n_i}\sum_{m=1}^{n_i} I_{1,i}(l) I_{1,i}(m)\right] + E\left[\sum_{l=1}^{n_j}\sum_{m=1}^{n_j} I_{1,j}(l) I_{1,j}(m)\right] - 2E\left[\sum_{l=1}^{n_i}\sum_{l=1}^{n_j} I_{1,i}(l) I_{1,j}(m)\right]\right).$$

Since beneficial mutations have identical $s$ irrespective of trait, the trait designation is a neutral marker. Conditioning inside the expectation on $n_i$ and $n_j$ results in random variables $\sum_{l=1}^{n_i} I_{1,i}(l)$ and $\sum_{m=1}^{n_j} I_{1,j}(l)$ that have binomial distributions with parameters $n_i$ and $n_j$, respectively, each with probability $p = U_1/(U_1 + U_2)$. This fact allows us to rewrite the expectations above as

$$\sigma_1^2 = \frac{s^2}{2}\left(p^2 E\left[n_i^2 + \left(\frac{1-p}{p}\right) n_i\right] + p^2 E\left[n_j^2 + \left(\frac{1-p}{p}\right) n_j\right] - 2p^2 E\left[n_i n_j\right]\right)$$

$$= p^2\left(\frac{s^2}{2} E[(n_i - n_j)^2] + \frac{s^2}{2}\frac{1-p}{p} E[n_i + n_j]\right),$$

As in Gomez et al. [3], we see that $\frac{s^2}{2} E[(n_i - n_j)^2] = \sigma^2 \approx v(U_1 + U_2, s, N)$ by Fisher's Fundamental Theorem, where $v(U, s, N)$ is Desai and Fisher's [1] result, given as Eq. (3) in our main text. $E[n_i + n_j] = \Pi$ is the average total pairwise heterozygosity at positively selected sites for two randomly selected individuals. Equation 30 in Desai et al. [4] provides an expression for $\Pi$, derived from the fitness-class coalescent of [5]. Specifically, we have

$$\Pi = \frac{2v}{s^2}\left(\ln\left(\frac{s}{U_1 + U_2}\right) + \frac{s}{\sqrt{\pi v}}\right),$$

where $v = v(U_1 + U_2, N, s)$. Substituting in for these expressions above provides

$$\sigma_1^2 = \frac{s^2}{2}\left(p^2 E\left[n_i^2 + \left(\frac{1-p}{p}\right)n_i\right] + p^2 E\left[n_j^2 + \left(\frac{1-p}{p}\right)n_j\right] - 2p^2 E\left[n_i n_j\right]\right)$$

$$= p^2\left(\frac{s^2}{2}E[(n_i - n_j)^2] + \frac{s^2}{2}\frac{1-p}{p}E[n_i + n_j]\right)$$

$$= p^2\left(v + v\frac{1-p}{p}\left(\ln\left(\frac{s}{U_1+U_2}\right) + \frac{s}{\sqrt{\pi v}}\right)\right)$$

which then yields the equation

$$\sigma_1^2 = v \cdot p^2 + vp(1-p)\left(\ln\left(\frac{s}{U_1+U_2}\right) + \frac{s}{\sqrt{\pi v}}\right). \tag{S3.9}$$

Similar steps to those above can be used to show that

$$\sigma_2^2 = v \cdot (1-p)^2 + vp(1-p)\left(\ln\left(\frac{s}{U_1+U_2}\right) + \frac{s}{\sqrt{\pi v}}\right). \tag{S3.10}$$

by simply starting with indicator variables $I_{2,i}(l)$ and $I_{2,j}(m)$ that identify mutations occurring in trait two, and recognizing that beneficial mutations occur on the second trait with probability $(1-p) = U_2/(U_1+U_2)$. With $\sigma_1^2$ and $\sigma_2^2$ in hand, we can solve for the covariance $\sigma_{12}$ using the relationship $v = v_1 + v_2 = (\sigma_1^2 + \sigma_{12}) + (\sigma_2^2 + \sigma_{12})$. Thus, $\sigma_{12} = \frac{1}{2}(v - \sigma_1^2 - \sigma_2^2)$, and hence

$$\sigma_{12} = vp(1-p) - vp(1-p)\left(\ln\left(\frac{s}{U_1+U_2}\right) + \frac{s}{\sqrt{\pi v}}\right). \tag{S3.11}$$

Since $v_k = \sigma_k^2 + \sigma_{12}$ for $k = 1, 2$, it follows that

$$v_1 = vp = \frac{U_1}{U_1+U_2} \cdot v(U_1+U_2, s, N) \tag{S3.12}$$

and

$$v_2 = v(1-p) = \frac{U_2}{U_1+U_2} \cdot v(U_1+U_2, s, N). \tag{S3.13}$$

The rates of fitness increase described by Eq. (S3.12) and Eq. (S3.13) can be converted into rates of adaptive substitutions $R_k$ ($k = 1, 2$), given our scenario of identical $s$ in both traits. Specifically, we have

$$R_1 = \frac{U_1}{U_1+U_2} \cdot \frac{v(U_1+U_2, s, N)}{s} \tag{S3.14}$$

and

$$R_2 = \frac{U_2}{U_1+U_2} \cdot \frac{v(U_1+U_2, s, N)}{s}. \tag{S3.15}$$

These expressions can be used to illustrate how clonal interference between traits impacts the number of beneficial substitutions in each trait. In particular, if we let $\alpha = U_2/U_1$, then

$$R_1 = \frac{1}{1+\alpha} \cdot \frac{s\log\left(N^2 s U_1(1+\alpha))\right)}{\log^2(s/U_1(1+\alpha))}. \tag{S3.16}$$

after substituting in Desai and Fisher's Eq. (4) on the right-hand side. The dependence of Eq. (S3.16) on $\alpha$ is depicted in figure S1.

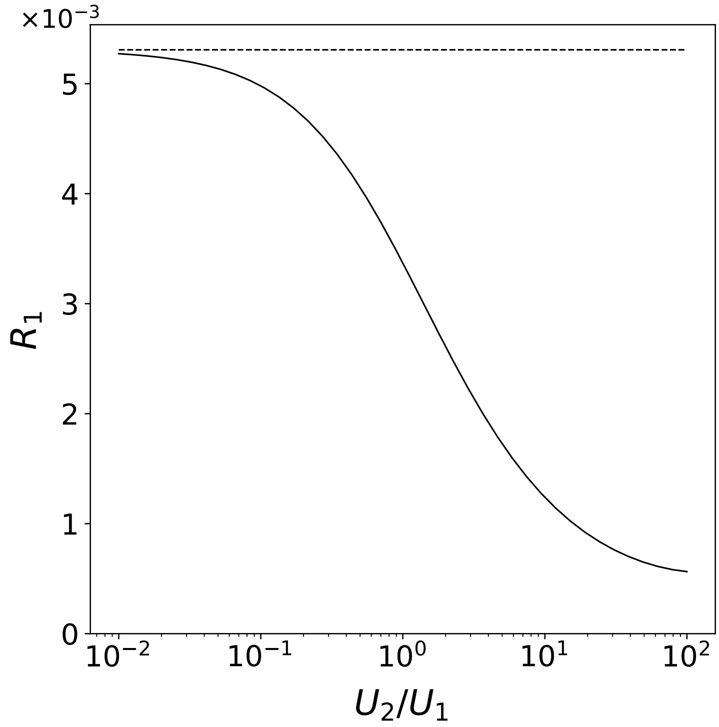


Figure S1: Solid line gives substitution rate of beneficial mutations in trait one as the ratio $U_2/U_1$, denoted $\alpha$ in Eq. (S3.16), increases. Dashed line provides substitution rate of beneficial mutations in absence of clonal interference with second trait. The parameters $N = 10^9$, $s = 0.01$, and $U_1 = 10^{-5}$ were used to generate the graph.
